# Arctic defaunation initiated a cascade of mammal–plant interaction shifts through dispersal dynamics

**DOI:** 10.1101/2025.06.23.661215

**Authors:** Sisi Liu, Kathleen R. Stoof-Leichsenring, Marc-Thorsten Hütt, Weihan Jia, Boris K. Biskaborn, Bernhard Diekmann, Darrell S. Kaufman, Hanno Meyer, Martin Melles, Luidmila A. Pestryakova, Ulrike Herzschuh

## Abstract

Amid accelerating defaunation, it remains unclear whether the loss of large mammals can trigger ecosystem collapse and through which functional pathways. Using sedimentary ancient metagenomics, we identify an overlooked role of large mammal defaunation in helping drive the collapse of the glacial mammoth-steppe. Network analyses identified rewiring of mammal interactions during the late glacial supporting cascading effects. Crucially, megafaunal-mediated long-distance seed dispersal buffered vegetation turnover during periods of rapid climate warming but weakened with progressive losses of vital dispersal interactions leading to the steppe–tundra’s transition to open woodland in the early Holocene. Trophic interactions did not offset turnover and became dominant afterwards with the more stable climate. We identified resilient interactions that may support recovery, suggesting the potential of rewilding to restore Arctic ecosystems.

**One-sentence abstract:** Ancient metagenomics reveals disrupted long-distance seed dispersal helped drive mammoth-steppe collapse and informs Arctic rewilding efforts.

## Introduction

Climate change is accelerating global declines, extinctions, and local extirpations of terrestrial animals (*1*), with large-bodied mammals—particularly herbivores and predators— being most at risk (*2*). Such declines can cause frequent loss of interactions, leading to functional extinction even while species persist (*3*). The extent to which these declines disrupt trophic (e.g., predation, herbivory) and mutualistic (e.g., seed dispersal) interactions, and how such disruptions unfold over time, remains insufficiently understood (*4, 5*). Yet, defaunation is not a uniquely contemporary phenomenon (*6*). A similar pronounced event happened around the end of the last glacial period (*7*), when many large-bodied mammals like mammoths declined and contracted in range. This coincided with the collapse of the vast mammoth-steppe ecosystem (*8*), once the largest biome on Earth.

While drivers of megafaunal extinction have been debated for decades, the ecological consequences—particularly shifts in biotic interactions—are only beginning to be understood (*9*). Crucially, Holarctic megafaunal declines unfolded gradually (*7*), likely altering terrestrial ecosystems well before their final extinction (*10*). This raises the hypothesis—so far untested— that cascading losses of biotic interactions due to defaunation may have contributed to the ecological reorganization of the mammoth-steppe ecosystem.

Beyond biotic interaction loss, the role of defaunation in enabling the emergence of novel interactions has received even less attention (*11, 12*). Different types of reconstructed interaction networks offer complementary insights: predator-prey webs, for example, suggest greater functional redundancy in the Pleistocene than Holocene, with release from top-predator loss leading to increases in megaherbivore density and distribution (*13*). Herbivory-plant interactions indicate that some surviving small-bodied herbivores shifted from browsing to mixed feeding between the Pleistocene and Holocene (*14*), potentially forming new trophic interactions. Plant–frugivore seed dispersal interactions exhibited profound rearrangement rather than complete structural collapse after late Pleistocene megafaunal loss (*11*). While this suggests a degree of functional replacement, it raises open questions about whether such interaction rewiring can buffer or instead contribute to shifts in plant community composition.

Additionally, the mechanisms behind the re-establishment of lost interactions—so-called “resilient links”—are still not well understood. Clarifying the role of functional traits in this process could provide important insights for coextinction risk assessments, particularly by accounting for interaction rewiring, the ability of a taxon to switch partners in response to perturbations as a potential buffer against cascading extinctions (*15*).

While much has been learned about past changes in taxonomic abundance across space and time, the dynamics of ecological interactions remain underexplored, as interactions are rarely preserved due to the sparse fossil record. Sedimentary ancient DNA (*sed*aDNA) is a powerful tool to infer biotic interactions by providing spatially and temporally extensive evidence of past taxonomic co-occurrence (*16, 17*). When combined with species traits, *sed*aDNA can help infer potential ecological interaction pathways with greater resolution than fossil records and more empirical grounding than trait-matching models alone.

To address these knowledge gaps, we combined two complementary datasets from Siberia and Alaska (Fig. 1A). One comprises 1-ka mammal occupancy and pollen records to resolve temporal ordering; the other, 5-ka metagenomic data to reconstruct mammal–plant interaction networks. Together, they allowed us to test whether mammal rewiring preceded vegetation reorganization and through which functional pathways it unfolded.

**Fig. 1.**
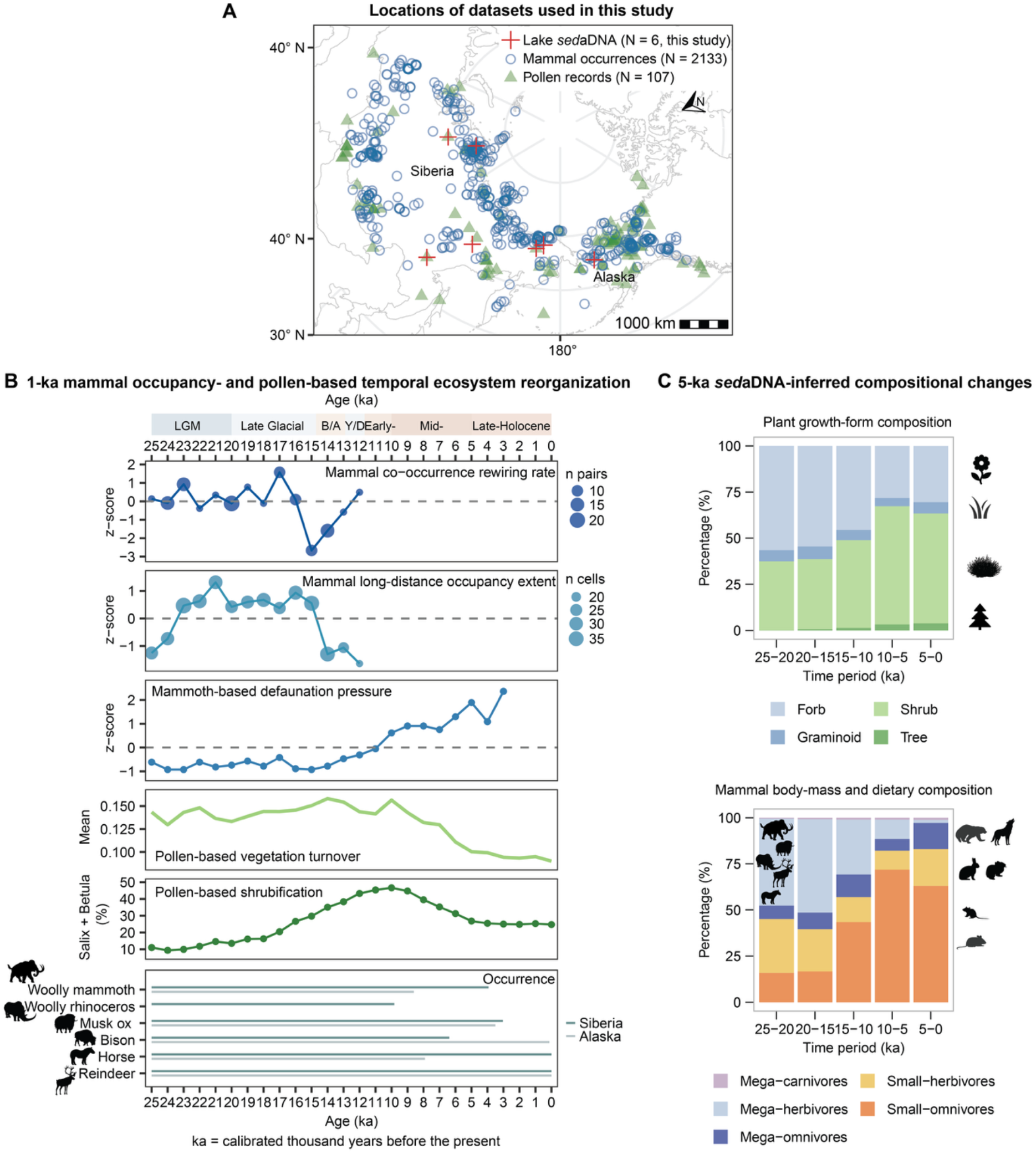
Fine-resolution occupancy and pollen records identify past-25-ka ecosystem reorganization mirrored by coarser *sed*aDNA-inferred compositional shifts. (**A**) Locations of the datasets used in this study across Siberia and Alaska. (**B**) 1-ka mammal occupancy and pollen records show temporal ecosystem reorganization from 25 to 0 ka. Mammal co-occurrence rewiring rate, mammal long-distance occupancy extent, and mammoth-based defaunation pressure summarize changes in mammal spatial overlap, broad-scale spatial occupancy, and mammoth spatial contraction, respectively. Pollen records show vegetation turnover and shrubification, and occurrence bars indicate the temporal presence of selected large herbivores. Top bar shading schematically indicates the relative warm–cold character of the major climatic intervals and is shown for visual context. (**C**) Upscaled to 5-ka intervals, *sed*aDNA-inferred composition captures the major ecological changes identified in the 1-ka records. Vegetation shifts from steppe-tundra to open woodland ecosystems. Concurrent shifts in mammal communities, from mega-herbivore-dominated to mega-omnivores with smaller-bodied mammals, indicate collapse of the mammoth steppe ecosystem. Percentages were calculated from 999 resampled community sets, with each lake contributing at most one sample per time period when available. Silhouettes were obtained from phylopic (https://www.phylopic.org/) under the CC0 1.0 Universal license.

### Millennial-scale ecosystem reorganization is robustly reconstructed from occupancy, pollen, and sedimentary ancient metagenomics

We compiled and harmonized 1-ka occupancy records for 10 well-sampled extinct and extant mammal taxa, integrating 1,273 directly dated megafaunal remains, 227 Siberian human occupation records, and ancient metagenomic datasets (table S1). Aggregated into 1-ka occupancy maps, these records yielded three spatial indices based on occupied grid cells and taxon overlap: co-occurrence rewiring, long-distance occupancy extent, and mammoth-based defaunation pressure (Materials and Methods; Fig. 1B and figs. S1–S2). These indices respectively quantify temporal reorganization of mammal spatial overlap, broad-scale spatial extent of occupancy, and contraction of the mammoth spatial footprint. We find that co-occurrence rewiring declined by 17–15 ka, long-distance occupancy extent contracted by 15– 12 ka, and mammoth-based defaunation pressure became pronounced mainly after 11 ka. Pollen-based vegetation turnover and shrubification, inferred from 107 pollen records (*18*), were strongest between 14 and 10 ka, marking intensified plant-community reorganization (*19*). Together, these records place mammal spatial restructuring before the strongest vegetation changes and mammoth-based defaunation signal, with the final regional losses of several large herbivores occurring afterwards (Fig. 1B).

Our six lake *sed*aDNA metagenomic (shotgun-sequencing) datasets yielded 80,869,509 reads for terrestrial ecosystem reconstruction, corresponding to 16 mammalian and 51 seed-bearing plant families (data S1). These reads, with a best identity of ≥95% against the comprehensive taxonomic reference database, were retained as part of a multi-step strategy (*20*) to minimize false positives and noise (Methods and Materials). Ancient origin was supported by read-length distributions (fig. S3) and terminal C-to-T substitutions across taxa and time periods (fig. S4), with representative coverage of mammals and plants (fig. S5). Concordance with metabarcoding data from the same cores supports family-level vegetation inferences from shotgun data (table S2). Aggregated into 5-ka time intervals, our shotgun data captured a broad transition detected in the independent 1-ka records: mammal assemblages are megaherbivore-dominated before ∼15 ka, whereas shrubs increase as megaherbivores decline after ∼15 ka (Fig. 1C and fig. S6). This concordance justifies using 5-ka *sed*aDNA networks for downstream analyses of interaction rewiring and compositional turnover, despite coarser temporal binning.

### Cascading effects: mammalian interactions shaped ecological turnover

We find that mammal co-occurrence rewiring led to pollen-based vegetation turnover by 2 ka, improving prediction beyond temperature change alone (ΔAIC = -4.9, *P* = 0.022, table S3; Fig. 2A). This sequence was inferred from time-lag models applied to the 1-ka mammal occupancy and pollen records, which tested mammal-leading, plant-leading, and climate-covarying alternatives. Climate changes were included because temperature and precipitation shifts are known to influence both Arctic vegetation turnover and megafaunal restructuring (*8, 21*), and were associated with the 1-ka records: temperature with vegetation turnover (ΔAIC = -10.9, *P* = 0.041, table S4) and precipitation with mammal rewiring (ΔAIC = -6.2, *P* = 0.028, table S5). Reverse-lag plant-leading models were unsupported (Fig. 2A and table S6), strengthening the directional interpretation of the mammal-leading sequence.

**Fig. 2.**
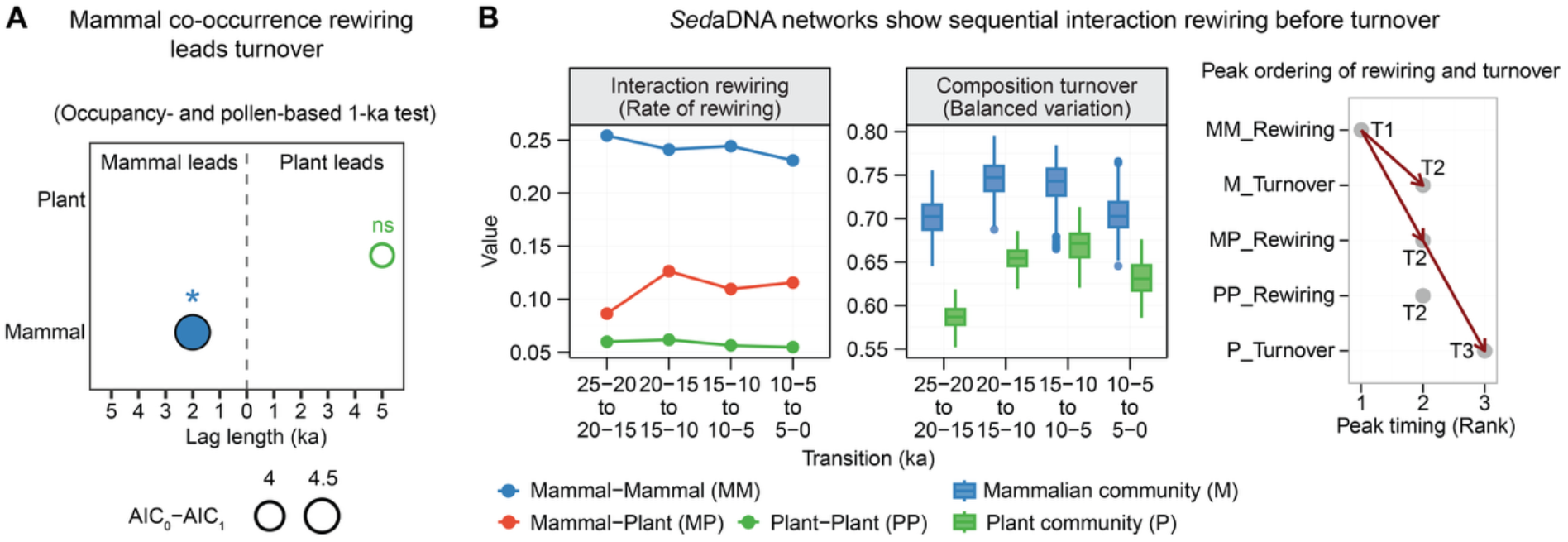
Mammal co-occurrence and interaction rewiring preceded plant turnover. (A) Mammal co-occurrence rewiring leads to pollen-based vegetation turnover by 2 ka in 1-ka lag models, whereas the reverse plant-leading model is not significant. Circle size is proportional to model improvement over the climate-only baseline (|AIC_0_–AIC_1_|). Significance codes are based on F tests with 999 permutations: * *P* < 0.05, and ns *P* ≥ 0.05. (B) At 5-ka resolution, *sed*aDNA networks show sequential mammal interaction rewiring before community turnover.

To address the challenge of inferring past ecological interactions from *sed*aDNA (*22*), we developed a validation pipeline for ancient ecological networks (Materials and Methods; figs. S7–S14; tables S7–S11). We then reconstructed ecological networks, comprising mammal– mammal, mammal–plant, and plant–plant interactions. We quantified and normalized interaction rewiring (i.e., changes in interactions among shared taxa, fig. S15), while compositional turnover was assessed separately. Specifically, a high rate of rewiring suggests rapid turnover in interaction structure, reflecting dynamic shifts in ecological roles or partner availability (*23, 24*).

Consistent with the 1-ka records, the 5-ka ecological networks show a mammal-first ordering of change (Fig. 2B). Mammal–mammal interaction rewiring peaks first (25–20 to 20– 15 ka), followed by delayed peaks in mammalian compositional turnover and mammal–plant and plant–plant interaction rewiring (20–15 to 15–10 ka). Mammal–plant interaction rewiring then precedes maximum plant composition turnover (15–10 to 10–5 ka).

Our network analyses reveal that mammalian interaction structure reorganized before mammal community composition changed. The early peak in mammal-mammal interaction rewiring (25–20 to 20–15 ka) likely reflects high interaction reshuffling potential under broad spatial overlap and low defaunation pressure (Fig. 1B). Megafauna was prominent, their large ranges and mobility (fig. S11) could have enabled rapid reorganization of interaction partners across regions while overall mammal composition remained comparatively stable. From ∼15 ka onward, the relative abundance of megaherbivores declined sharply (Fig. 1C), coinciding with a slight decline in mammalian interaction rewiring from 20–15 to 15–10 ka. A concurrent rise in network modularity (fig. S16) suggests that, rather than forming well-connected webs, surviving mammals increasingly clustered into more isolated groups with fewer cross-group links. Reduced interaction redundancy likely increased vulnerability to perturbation, contributing to the pronounced turnover in composition during this transition. Thereafter, mammalian interaction rewiring remained low and compositional turnover declined into the late Holocene (5–0 ka), consistent with community simplification and reduced resilience (*14*).

Neither the 1-ka nor the 5-ka dataset shows a peak in plant compositional turnover before ∼15 ka, when megaherbivores remained abundant. This finding suggests that trophic effects alone did not drive vegetation shifts, challenging the commonly held view that strong grazing pressure necessarily produces pronounced vegetation change, including reduced tree cover (*25*) and increased abundance of grazing-tolerant grasses and forbs (*26, 27*). One explanation is that inferences drawn from already defaunated ecosystems may overemphasize local trophic forcing (*28*).

Instead, our results identify long-distance spatial connectivity as a stabilizing mechanism at the landscape scale. In the 1-ka dataset, vegetation turnover increased with mammal co-occurrence rewiring, but the increase was weaker where mammals occupied a larger spatial extent (*P* = 0.015, fig. S17). This effect remained after accounting for temperature (*P* = 0.002, table S12).

Furthermore, our 5-ka *sed*aDNA networks identified megafauna-linked seed dispersal as a key mechanism by which mammal spatial connectivity stabilized vegetation change. This mechanism was inferred from pathway prediction and quantified using edge betweenness (EB) centrality for mammal–plant interactions (Materials and Methods; figs. S18 and S19; tables S13–S16). High EB values identify links most important for maintaining ecological connectivity (*29, 30*). Dispersal-linked connectivity eroded progressively, with the largest net loss from 20-15 to 15-10 ka (Fig. 3A; fig. S20). This loss was concentrated in megafauna-associated dispersal links (Fig. 3B), indicating that megafaunal seed dispersal helped maintain ecological connectivity (Supplementary Text1).

**Fig. 3.**
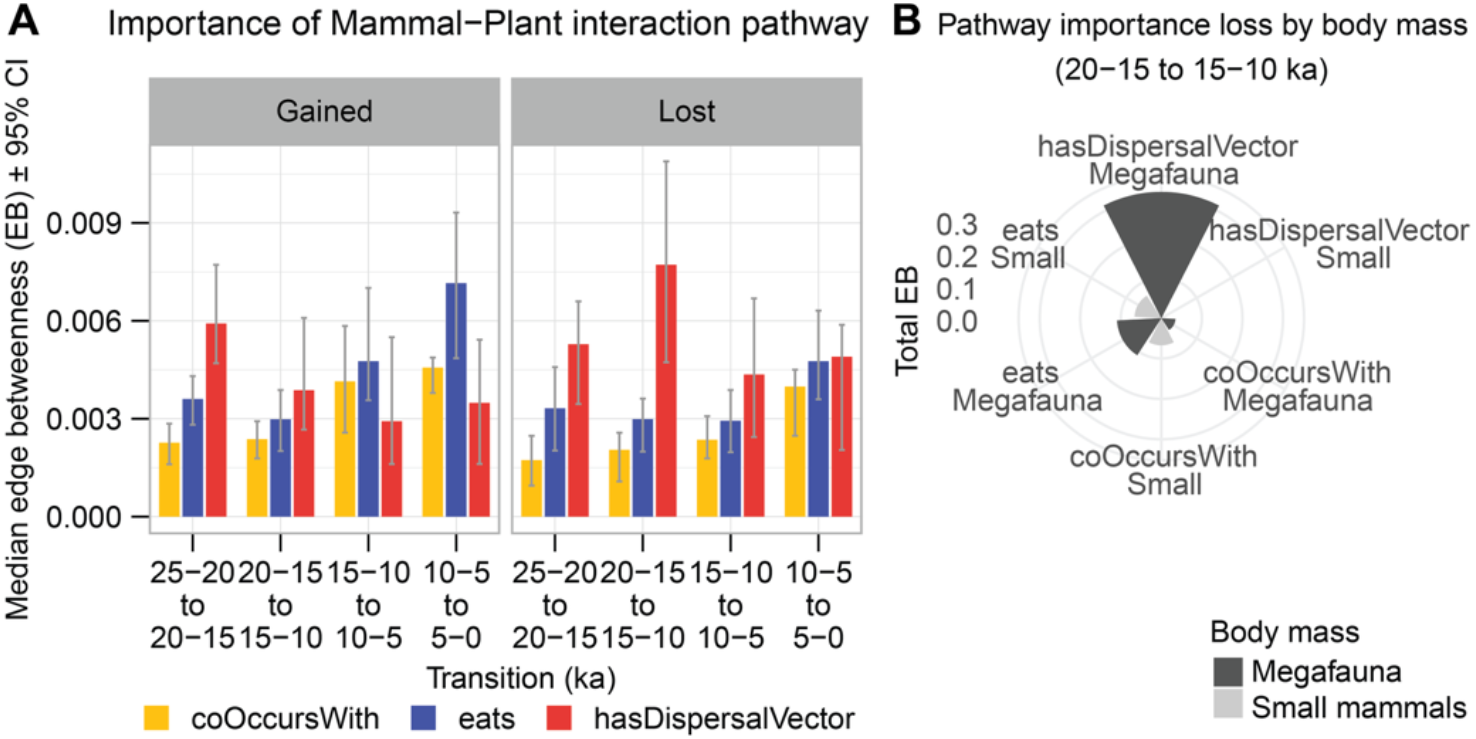
Dispersal-related connectivity eroded as trophic links gained prominence over the past 25 ka. (**A**) Median weighted edge betweenness (EB) of gained and lost mammal–plant links across four transitions, grouped into habitat co-occurrence (coOccursWith), trophic (eats), and seed-dispersal (hasDispersalVector) pathways. EB was weighted by rank-scaled relative abundance of linked mammal and plant taxa to approximate interaction influence on ecosystem-level connectivity. Error bars show 95% bootstrap confidence intervals. (**B**) Total weighted EB of links lost during the main collapse transition (20–15 to 15–10 ka), grouped by pathway and mammal body-mass class, showing that megafauna account for most connectivity loss in dispersal interactions. Pathways were assigned using the Global Biotic Interactions (GloBI) database, and body-mass classes were based on Bayesian body-mass estimates (fig. S11). Only links with robust pathway predictions under trait uncertainty were included in the main pathway analyses.

From 15-10 ka onward, trophic links, and secondarily habitat co-occurrence links, gained relative importance (Fig. 3A). Small mammals contributed substantially to these gains (fig. S21), with the strongest late-Holocene increase concentrated in trophic links, while recovery of dispersal links remained limited (fig. S22 and Supplementary Text2). Loss of long-distance seed-dispersal connectivity likely reduced the movement and deposition of seeds into suitable habitats as environments changed, increasing vulnerability to plant turnover and biome-scale reorganization. Consistent with this mechanism, dispersal-network erosion preceded peak plant compositional turnover (Fig. 2B) and may have contributed to steppe-tundra collapse.

More broadly, regulation of steppe-tundra appears to have shifted from dispersal-based integration to a more local, patch-dependent trophic regime. As shrubification progressed and steppe-tundra transitioned toward shrub-tundra, reduced long-distance seed dispersal would have disadvantaged strict grazers, favored mixed feeders and browsers, and increased extinction risk among open-habitat taxa (*19, 21, 31*). Megafaunal extinction thus appears as a late-stage expression of an earlier functional transition (Fig. 4). This temporal ordering is statistically consistent with a mammal-initiated interaction cascade (one-sided exact permutation test, *P* = 0.01; all 5,040 permutations).

**Fig. 4.**
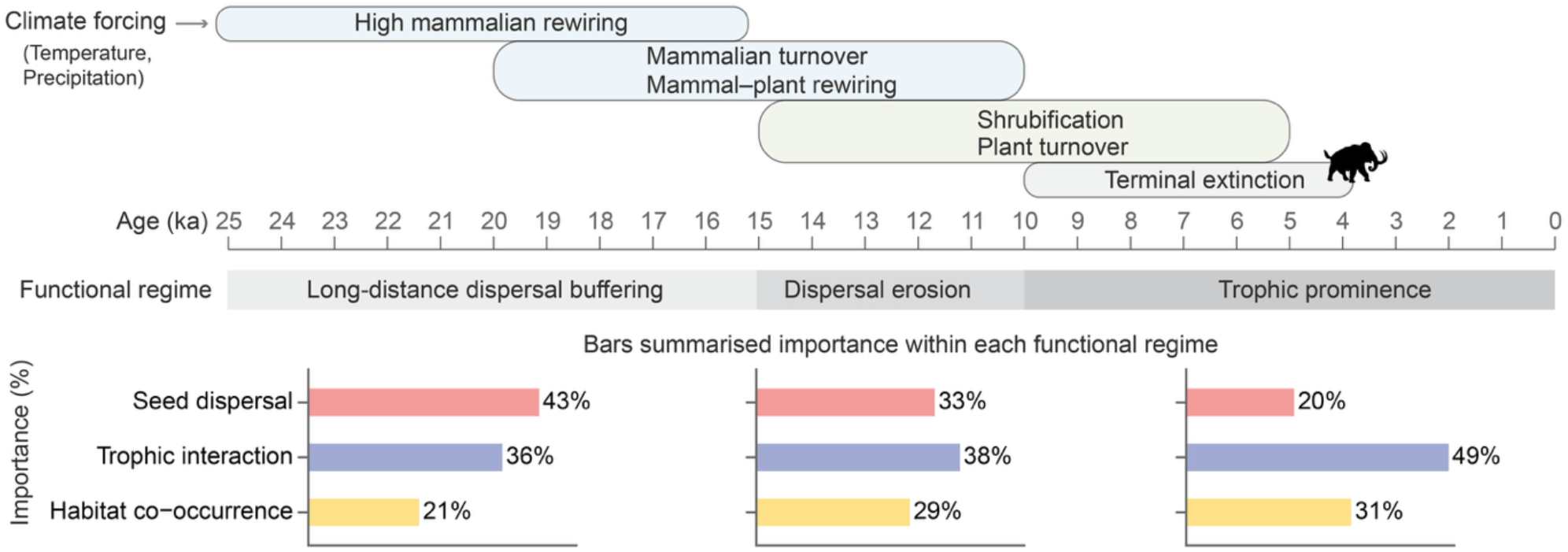
Summary of the mammal-initiated cascade. The upper bars summarize the relative timing of major climate-associated mammalian and plant-community events across the past 25 ka (integrating 1-ka and 5-ka evidence). The middle band shows the inferred functional regime shift from long-distance dispersal buffering of ecological connectivity to dispersal erosion and later trophic prominence. The lower bar plots show the relative connectivity contributions of seed-dispersal (hasDispersalVector), trophic (eats), and habitat co-occurrence (coOccursWith) pathways across these three regimes, calculated from links with robust pathway predictions under trait uncertainty. Formal permutation-based ordering tests were conducted on 1-ka events only (see Materials and Methods).

Cumulatively, our findings reveal a previously overlooked mechanism: seed dispersal interactions played a pivotal role in dampening vegetation change—long-distance megafaunal movement helped delay plant community turnover by millennia, even after mammalian community turnover had peaked.

### Key mammal–plant interactions and their ecosystem-level influence

We find that key seed dispersal interactions are strongly associated with megaherbivores (e.g., Elephantidae, Bovidae, and Equidae) and are predominantly positive (Fig. 5A–E). This suggests their important role as long-distance dispersers, maintaining ecological connectivity by linking otherwise distant or isolated vegetation patches (*32*). In contrast, key negative trophic interactions are concentrated in mega-omnivores such as Ursidae (Fig. 5A–B and D–E), likely reflecting an extractive foraging strategy shaped by low site fidelity and dietary opportunism (*33*). This pattern of foraging may have limited opportunities for mutualistic interactions. Moreover, cervids, though also classified as megaherbivores, primarily contributed to positive trophic interactions. This suggests that they functioned more as localized consumers than as landscape-scale seed vectors, consistent with their relatively short dispersal distances and generalist herbivory (fig. S11). These findings highlight that megafauna—even within herbivores—are functionally diverse.

**Fig. 5.**
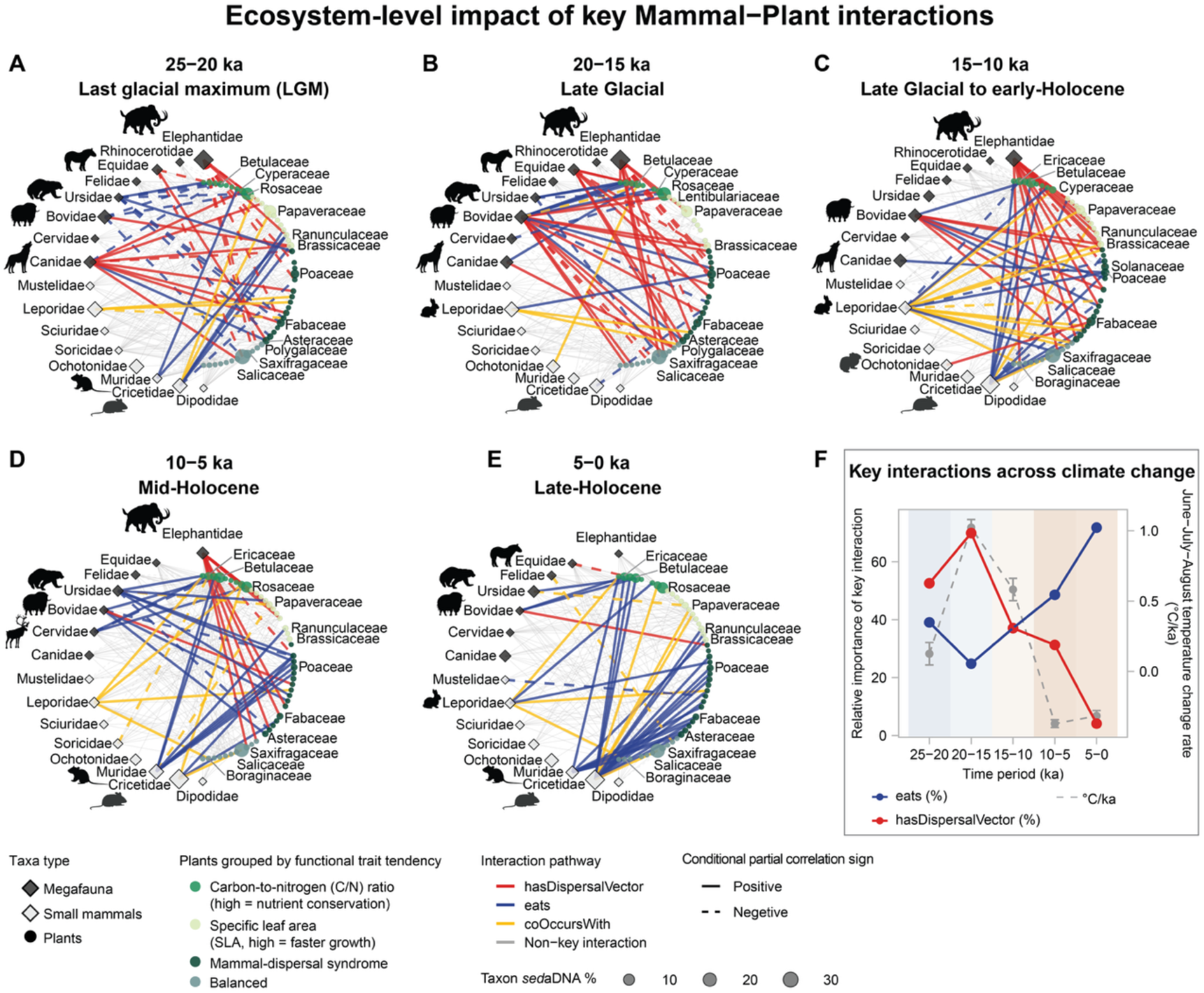
Temporal dynamics of key mammal–plant interactions at the ecosystem-level. (**A–E**) Reconstructed mammal–plant networks. Node size reflects sedaDNA-based relative abundance; shapes distinguish megafauna, small mammals, and plants. Mammals are ordered by decreasing dispersal distance, plants by multivariate trait similarity (table S17). Plant labels show the three most abundant families within each trait group. Link colors indicate interaction pathways. Links with weighted EB above the third quartile (Q3) are highlighted as key interactions (thicker lines); remaining robust links are shown in the background. All displayed links were robust to trait uncertainty. Larger versions with full plant labels are provided in fig. S23. Silhouettes were obtained from phylopic (https://www.phylopic.org/) under the CC0 1.0 Universal license. (**F**) Relative pathway importance through time, expressed as the percentage of total weighted EB among robust key links, shown alongside the June–July–August temperature change rate. Background shading shows the relative warm–cold character of each time period.

Across Siberia and Alaska, bottom-up frameworks attribute late-glacial to early Holocene ecosystem turnover to climatic reorganization that degraded the mammoth steppe as a productive forage system, with warming and wetting favoring shrub- and peat-dominated mosaics over graminoid-rich habitats (*21, 31, 34*–*36*). Our results identify a network-state mechanism that shaped how climate-driven bottom-up forcing affected plant communities. Key dispersal links increased in importance from 25–20 to 20–15 ka as temperatures rose (Fig. 5F). This strengthened dispersal connectivity likely maintained seed movement among functionally distinct plant groups, including graminoids, forbs, and shrub-associated taxa (Fig. 5A–B). As these dispersal links became dominant cross-community connectors (fig. S24), they may have helped these groups track post-LGM warming and limited strong vegetation turnover. The critical transition occurred when defaunation eroded mammals carrying key dispersal interactions during 15–10 ka (Fig. 5C), reducing long-distance seed movement and shifting regulation toward local trophic pathways. This reorganization suggests that climate effects on plant communities strengthened after ∼15 ka once long-distance seed dispersers declined.

After 15 ka, key trophic links gained relative importance, suggesting that the reassembled network became increasingly shaped by forage-mediated filtering. This aligns with Guthrie’s dietary hypothesis, which emphasizes the filtering of large grazers through changes in plant quality and defenses (*21, 35*). Continued shrubification likely intensified fragmentation, limited recolonization and habitat tracking, and reduced the re-linking of suitable habitat patches through seed dispersal during the Holocene. Together with the eventual loss of intercontinental dispersal routes, these changes likely made long-distance connectivity increasingly difficult to maintain. As graminoid-rich habitats declined, open-habitat specialists such as mammoths (Elephantidae) likely lost forage, whereas flexible browsers such as cervids (*19*), omnivores (*36*), and small mammals (*21*) were better able to persist. This selective loss of forage likely contributed to the staggered decline and eventual loss of open-habitat megafauna, while more flexible herbivores persisted to the present (Fig. 1B). Our findings suggest that modern Arctic shrubification may especially increase vegetation turnover in ecosystems already characterized by weakened mammal-mediated seed dispersal (*37*).

### Functional traits determine turnover and resilience in mammal–plant mutualisms

We find that positive mammal–plant interactions showed strong period specificity rather than broad resilience. Among inferred positive interactions, 27.1% were classified as glacial-specific, 26.1% as resilient, and 19.8% as Holocene-specific. This pattern indicates that many glacial-period interactions were not reassembled under warmer Holocene conditions. Generalized linear mixed-effects models showed that glacial-specific interactions tended to involve plants with higher carbon-to-nitrogen ratio values (Fig. 6A), with additional support for associations involving seed- and vertebrate-consuming mammals (table S18). This suggests that glacial positive associations were primarily linked to nutrient-conservative plants, with possible contributions from trophically flexible consumers under glacial resource constraints. Abundant megafaunal dispersers may have reduced dispersal limitation, promoting more diffuse plant–mammal coupling and greater flexibility in partner switching rather than reliance on specialized dispersal traits.

**Fig. 6.**
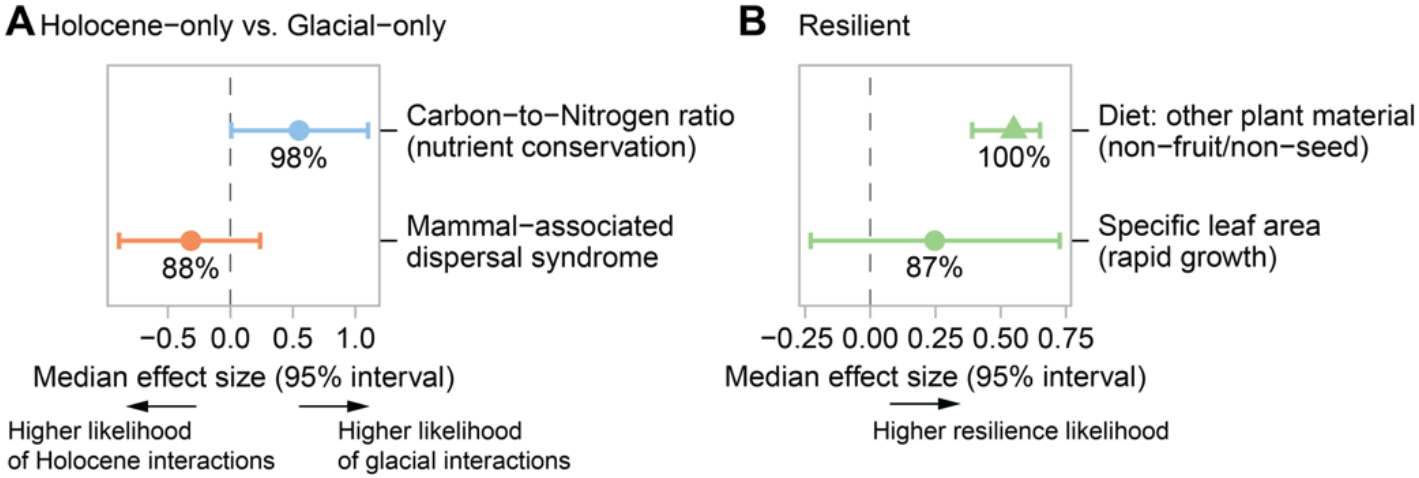
Trait effects on mammal–plant positive interactions over the past 25 ka. Median trait effects on (**A**) the likelihood of glacial-only, Holocene-only, and (**B**) resilient (i.e., lost and later re-established) positive interactions. Panels show the top two traits selected from 15 tested traits based on directional consistency and evidence tier. Points and horizontal bars indicate median effect sizes and 95% intervals accounting for family-level trait uncertainty. Percentages indicate the frequency of consistent effect direction under trait uncertainty. Models include mammal and plant identity as random effects.

In contrast, Holocene-specific positive interactions showed only a weak increase in mammal-associated dispersal traits (Fig. 6A), suggesting that reassembly did not yield a broadly mammal-dispersed flora. Notably, the Holocene dispersal network shifted toward larger-seeded plants (fig. S25), suggesting greater reliance of the remaining dispersal structure on taxa potentially dependent on mammal-mediated movement. The warmer and more stable Holocene likely made long-distance dispersal especially beneficial for those larger-seeded plants by improving opportunities for recruitment and range shifts (*38, 39*). At the same time, reduced availability of dispersers may have intensified partner dependency in these mutualistic relationships. This could have been further exacerbated by weaker wind regimes compared to the glacial period (*40*), which limited the effectiveness of wind dispersal and increased reliance on mammal-mediated seed dispersal. Thus, the loss of these commensalisms and mutualisms due to defaunation likely increased plant extinction that occurred during the early Holocene (*41*), while potentially leaving the remaining dispersal-mediated plants more vulnerable to further defaunation under continued warming (*42*).

Resilient interactions were primarily associated with mammals with broader, more flexible plant use, whereas plants with higher specific leaf area—a trait linked to rapid growth in disturbed environments—emerged as a secondary positive directional signal (Fig. 6B and table S19). This suggests that re-establishment of lost links depended more on the dietary plasticity of generalist consumers than on the persistence of specialized plant–animal partnerships. Together with modern evidence that mammal–plant interactions are strongly trait-dependent (*43, 44*), our results extend this trait-based perspective to millennial-scale interaction reassembly, highlighting mammals with broad dietary niches as key contributors to positive-link reassembly since the LGM.

## Conclusions

Our time-series *sed*aDNA data offer novel evidence that rewiring of mammalian interactions preceded ecological cascades unfolding over millennia, ultimately contributing to the collapse of the Arctic mammoth-steppe. We identified that mammal-associated seed dispersal was critical for maintaining ecosystem connectivity, but this role deteriorated as dispersers—particularly megaherbivores—were lost. These insights underscore the long-term, often delayed consequences of defaunation, implying that today’s human-driven losses may likewise trigger ecological shifts that unfold far beyond the current century.

Our findings refine the prevailing view that the disappearance of the mammoth-steppe biome is primarily due to climate warming that leads to vegetation change and, subsequently, to the loss of megafauna (*8*). Instead, our results highlight the active role of megafauna in shaping vegetation composition through long-distance seed dispersal, suggesting that restoring lost interaction functions could be an important consideration in rewilding under ongoing environmental change. Furthermore, we show that different megafauna contributed to ecosystem functioning via distinct interaction pathways with plants, highlighting those functional roles, not only species presence, may influence restoration outcomes.

Looking ahead, our findings suggest that interaction rewiring—detected before major compositional shifts—may serve as an early-warning signal of ecosystem reorganization. With finer temporal resolution, ancient metagenomic records could help identify subtle yet functionally significant changes, offering a quantitative framework to anticipate ecological tipping points.

## Acknowledgments

We used ChatGPT to improve language readability and clarity. All AI-generated outputs were carefully reviewed and edited by the authors. We thank Cathy Jenks for English proofreading and Janine Klimke for assistance with genetic laboratory work. We thank Christiane Böeckel for assistance with virus database indexing and integration into the NCBI taxonomy. We also thank Lars Harms for implementing the deduplication step in bioinformatics and for supporting software installation in the HPC environment.

## Author contributions

Conceptualization: S.L., U.H.

Methodology: S.L., K.R.S., U.H.

Validation: S.L., K.R.S, D.S.K.

Formal analysis: S.L

Investigation: K.R.S., W.J., D.S.K., S.L., U.H., B.K.B., B.D., H.M., M.M., L.A.P.

Resources: U.H., B.D., M.M., D.S.K.

Data curation: S.L., K.R.S., D.S.K., U.H.

Writing – original draft: S.L.

Writing – review & editing: S.L., U.H., M.T.H., D.S.K., K.R.S., B.K.B., M.M., B.D., W.J, H.M., L.A.P.

Visualization: S.L.

Supervision: U.H., S.L.

Project administration: U.H., S.L., K.R.S.

Funding acquisition: U.H., D.S.K., M.M.

## Data and materials availability

For transparency and reproducibility, source data and codes are informative in supplementary materials and/or deposited in open access platforms. Specifically, the raw metagenomics (shotgun sequencing) data are available in European Nucleotide Archive (ENA) with BioProject accession PRJEB94536 (Lake Levinson Lessing), PRJEB80877 (Lake Lama), PRJEB80635 (Lake Ilirney), PRJEB80642 (Lake Bolshoe Toko), PRJEB82635 (Lake Ulu), PRJEB82717 (Lake Salmon). The source codes for bioinformatics, taxonomic reference indexing, taxonomic classification, and ancient damage analysis are archived on Zenodo (https://zenodo.org/records/17974857). Source data, sequencing metadata, sample information, read counts, taxonomic classification matrices, environmental reconstructions, significant consensus networks, and predicted interaction pathways are provided in data S1–S17. The source code used for statistical analyses is provided in code S1–S31 and software S1. Rendered HTML tutorials are available in the GitHub repository (https://github.com/sisiliu-research/sedaDNA_ArcticCascadeEffect), which was included during the resubmission to assist peer reviewers with visualization and interpretation of the analyses.

## Supplementary Materials

Materials and Methods

Supplementary Text1

Supplementary Text2

Figs. S1 to S25

Tables S1 to S19

Data S1 to S17

Code S1 to S31

Software S1

References and Notes (*45*–*118*)

