## Supplemental Materials and Methods for "Arctic defaunation initiated a cascade of mammal–plant interaction shifts through dispersal dynamics"

#### Sites settings

The six lake sediment cores investigated span the Pleistocene–Holocene transition and are located across Siberia and Alaska, encompassing the Beringia region (Fig. 1A). This spatial distribution enables the investigation of mammalian and vegetation changes across areas formerly occupied by the Pleistocene mammoth-steppe.

The geophysical and vegetation characteristics of the catchments and surrounding areas of Lakes Levinson Lessing [74.47°N, 98.67°E; 48 m above sea level (a.s.l.)], Lama (69.53°N, 90.20°E; 53 m a.s.l.), Ilirney (67.35°N, 168.32°E; 1790 m a.s.l.), Ulu (63.34°N, 141.04°E; 950 m a.s.l.), and Bolshoe Toko (56.25°N, 130.50°E; 903 m a.s.l.) have been previously described in detail (*41*, *45–47*). In summary, these sites span a broad range of geological settings across western Russia and eastern Siberia, from rims of the Putorana Plateau (Lake Lama) and the Byrranga Mountains of the Taymyr Peninsula (Lake Levinson Lessing) to the Stanovoy Mountain Range (Lake Bolshoe Toko), Oymyakon Upland (Lake Ulu), and Chukotka near the Anadyr Mountains (Lake Ilirney). All sites experience a strongly continental climate, with mean January temperatures ranging from −65°C to −28.8°C and mean July temperatures between 12.1°C and 34°C. Bedrock geology is generally dominated by crystalline and metamorphic units, including granite, gneiss, and schist, while vegetation ranges from sparse typical tundra to *Larix*-dominated taiga (*41*, *45–47*).

Lake Salmon (64.91°N, 164.99°W; 208 m a.s.l.) is a glacially formed lake situated south of the Kigluaik Mountains, about 47 km north-northeast of Nome, Alaska. The lake covers 7.5 km², reaches a maximum depth of 21.6 m, and has a catchment area of 215 km². The catchment is primarily underlain by silicate-rich, high- to medium-grade metamorphic rocks of the Nome Group, including amphibolite- and granulite-facies of gneiss and biotite schist (*48*). Mean January and July temperatures are −14.6 °C and 10.5 °C, respectively, based on records from meteorological station Nome (49 km away, 11 m a.s.l.). The modern vegetation consists of dwarf to tall shrub tundra and dry-to-moist tundra communities, dominated by species such as *Alnus crispa*, *Calamagrostis canadensis*, *Spiraea beauverdiana*, *Senecio lugens*, *Betula nana*, *Loiseleuria procumbens*, *Stereocaulon paschale*, and *Rhododendron camtschaticum* (*49*).

### Cores, chronologies, and *sed*aDNA materials

Core collection, radiocarbon dating, and age–depth models have been previously described for Lakes Levinson Lessing (core ID: Co1401) (*50*), Lama (PG1341) (*45*), Ilirney (EN18208) (*51*), Ulu (EN21103) (*47*), Bolshoe Toko (PG2133) (*46*), and Salmon (EN22502) (*52*). We used the published age–depth models for all six cores. For transparency and comparability, we summarize below the dated materials, chronological controls, calibration approaches, treatment of potential old-carbon effects, and the relevance of chronological uncertainty to the 5-ka temporal bins used for the *sed*aDNA shotgun analyses.

Levinson-Lessing Lake, core Co1401. The published chronology for core Co1401 is based on a multi-control Bayesian age–depth model constrained by the modern sediment surface, palaeomagnetic tie points and excursions, Accelerator Mass Spectrometry (AMS) $^{14}$C dates, Optically Stimulated Luminescence (OSL) ages, and pollen-based comparison points. Radiocarbon dates were obtained from macro-organic remains and calibrated using the IntCal20 calibration curve. The original study considered possible $^{14}$C age overestimation caused by hard-water effects or redeposited organic material, but also noted that OSL ages may be affected by water-content correction, sediment compaction, or incomplete luminescence-signal resetting. The study therefore did not apply a single numerical old-carbon correction; instead, the Bayesian age–depth model integrated multiple chronological controls with their associated uncertainties. In the present study, we use only the 0–25 ka interval of this core. Within this interval, the reported calibrated $^{14}$C 2σ uncertainties are ≤0.61 ka.

Lama Lake, core PG1341. The published chronology for core PG1341 is described in supplement 3 of ref. (*45*). It is based mainly on radiocarbon dating of bulk sediment total organic carbon (TOC) samples, supplemented by dated woody remains. The published chronology applied a 4460-year old-carbon correction to the measured $^{14}$C ages before age–depth modelling. Thus, potential old-carbon effects were explicitly treated in the published chronology. The age–depth model was calibrated with IntCal20. Within the interval used in this study, the maximum modelled age uncertainty is ±0.94 ka. Within the 0–25 ka interval used here, the Lama Lake age–depth model had a mean 95% half-range uncertainty of 0.61 ka (median 0.60 ka; range 0.065–0.995 ka).

Ilirney Lake, core EN18208. The published EN18208 chronology is based on AMS $^{14}$C dating of bulk-sediment TOC and OSL dating, because macrofossils were not available. A surface-sediment age of 1,721 ± 28 $^{14}$C yr BP was used to estimate an old-carbon effect, which was subtracted from down-core $^{14}$C ages before IntCal20 calibration. The age–depth model was constructed in Undatable using four OSL ages and 17 reservoir-corrected $^{14}$C dates. Dates between 282 and 755 cm, corresponding to ~25.9–35.6 cal ka BP, were excluded because of $^{14}$C–OSL discrepancies; this interval is older than the Ilirney shotgun record used here, which extends only to 19.8 cal ka BP. Accordingly, the Ilirney age–depth model had a mean 95.4% half-range uncertainty of 0.96 ka (median 0.78 ka; range 0.091–1.92 ka).

Ulu Lake, core EN21103. The published EN21103 chronology is based on AMS $^{14}$C dating of bulk-sediment TOC and a Bayesian Bacon age–depth model calibrated with IntCal20. A reservoir effect of 351 ± 24 $^{14}$C yr, estimated from a surface sample, was subtracted from all $^{14}$C ages before calibration; age reversals from deep in the core were excluded or selectively used in the published model. In this study, we used Ulu samples back only to 24.098 cal ka BP; within this interval, the age–depth model had a mean 95% half-range uncertainty of 0.51 ka (median 0.48 ka; range 0.114–1.51 ka).

Bolshoe Toko Lake, core PG2133. The published PG2133 chronology is based mainly on AMS $^{14}$C dating of bulk sediment, with woody plant remains incorporated into the age model. The Holocene part of PG2133 was correlated with the higher-resolution core PG2208 from the same lake using Zr counts and total carbon content, and tie points between the cores were used to transfer age information to PG2133. The age–depth model was calibrated using IntCal20. Potential old-carbon effects were addressed by applying a 1,203-yr reservoir correction and excluding dates interpreted as affected by old organic carbon. Within the 0–25 ka interval, the Bolshoe Toko age–depth model had a mean 95% half-range uncertainty of 1.10 ka (median 1.11 ka; range 0.048–2.06 ka).

Salmon Lake, core EN22502. The published EN22502 chronology is based on AMS $^{14}$C dating of 49 macrofossil samples extracted from the sediment core and combined with short-lived isotope constraints. The dated materials consist mainly of plant macrofossils, including moss remains, vascular plant leaves/stems, woody fragments, and unidentified plant remains, with occasional chitinous or invertebrate remains. Radiocarbon ages were measured at the Arizona Climate and Ecosystems Isotope Laboratory, pretreated using weak acid, and the age–depth model was calibrated using IntCal20. Because the chronology is based primarily on macrofossils rather than bulk sediment, the risk of old-carbon contamination is reduced; no additional old-carbon correction was applied in the present study. Within the ≤25 ka interval used here, Salmon Lake age-model uncertainty averaged 0.35 ka as the 95% half-range, with values ranging from 0.009 to 0.77 ka.

Taken together, the reported radiocarbon and modelled age uncertainties for the intervals used in the sedaDNA shotgun analyses are substantially smaller than the 5-ka temporal bins, supporting the use of the published chronologies for multi-millennial comparisons and interpretations.

In addition to chronological considerations, core opening and subsampling were conducted in a dedicated laboratory (approx. 4–10 °C) at the GFZ Helmholtz Centre for Geosciences in Potsdam, a facility where no molecular genetic work is performed. All cleaning procedures followed established protocols to avoid contamination from modern DNA and cross-contamination between sediment samples (*53*). To reduce contamination risk, the exposed core surface was removed using a sterile, single-use blade. Only the inner sediments were stored in 8 mL sterile tubes at −20 °C until DNA extraction at the Alfred Wegener Institute, Helmholtz Centre for Polar and Marine Research (AWI) in Potsdam.

### DNA extraction, library preparation and shotgun sequencing

Molecular genetic analyses of sediment samples were carried out in the dedicated palaeogenetic laboratories at AWI, Potsdam, Germany. DNA was extracted from all lake sediment cores following a uniform protocol established for the study (*41*, *45–47*). In brief, each DNA extraction batch included nine sediment samples and one extraction control. Extractions followed a partially modified protocol of the PowerMax® Soil DNA Isolation Kit (QIAGEN, Germany) (*54*). The initial DNA concentration of the sedimentary ancient DNA (sedaDNA) isolates and blanks was measured using 1 µL of extract with the dsDNA BR Assay and Qubit® 2.0 fluorometer (Invitrogen, USA). Extracts were then concentrated and purified using the GeneJET PCR Purification Kit (Thermo Fisher Scientific Inc., USA), followed by a second concentration measurement. All purified sedaDNA isolates and blanks were diluted to 3 ng/µL, aliquoted into 15 µL portions to avoid repeated freeze–thaw cycles, and stored at −20 °C as working stocks for library preparation.

DNA libraries for all six sediment cores were prepared using a single-stranded DNA library protocol (*55*), including the second adapter ligation (CL53/CL72) performed on a thermomixer (*56*). Full details are described for Lake Lama (*57*) and were followed for other five cores. In this study, 29 (Levinson Lessing), 17 (Ilirney), 24 (Ulu), 27 (Bolshoe Toko), and 29 (Salmon Lake) *sed*aDNA isolates were processed for library preparation. Each library batch included 30 ng of DNA extract, at least one extraction blank, and one library blank. All libraries were quantified using quantitative polymerase chain reaction (qPCR), indexed with P5 (01–06, 11, 15–18, 46, 51, 64, 67, 72) and P7 primers (20, 21, 26, 27, 29, 40, 57, 60, 65, 82, 88, 91–96), purified using the MinElute PCR Purification Kit (Qiagen, Germany), and initially quality-checked on the 4200 TapeStation system (Agilent, G2991AA). Library-specific sequencing information is summarized in data S2. In total, 168 *sed*aDNA samples including 42 previously published shotgun-sequenced samples from Lake Lama (*57*), and 69 controls were processed for bioinformatic analysis.

### Bioinformatic analyses

We assessed the quality of raw paired-end reads (10,235,808,874) using FastQC (v0.11.9) (*58*). Duplicate reads were first removed using clumpify.sh from BBMap (v39.01, dedupe=t, https://github.com/bbushnell/BBTools/blob/master/clumpify.sh) (*59*). Adapter trimming, quality filtering, error correction, and merging of paired-end reads were then conducted using fastp (v0.23.2) (*60*), applying the same parameter settings as described by Liu et al. (*17*). Specifically, these steps included removal of low-complexity reads, length filtering, trimming of polyG and polyX tails, and read merging requiring a minimum overlap length of 30 bp, a maximum mismatch limit of 5, and a maximum mismatch percentage of 20%. The resulting merged reads were subsequently deduplicated using dedupe.sh from BBMap (v39.01, https://github.com/bbushnell/BBTools/blob/master/dedupe.sh) (*59*) to further remove redundant sequences. The quality of the merged reads was subsequently assessed using the FastQC (v0.11.9) (*58*).

We performed taxonomic classification on merged reads (5,755,136,011) via end-to-end alignment with Bowtie2 (v2.5.1) (*61*). Classification was conducted using ngsLCA (*62*), as implemented in MetaDMG (v0.37) (*63*), which subsequently assessed postmortem damage patterns to evaluate the ancient origin of the classified taxa. To improve accuracy and sensitivity of taxonomic classification, we established a curated reference database incorporating all available RefSeq genomes from National Center for Biotechnology Information (NCBI, as of August 14, 2023), the full NCBI nucleotide (NT) database (as of September 19, 2022), chloroplast and mitochondria genomes of plant species occurring in SibAla (Siberia-Alaska) region [as defined in ref. (*41*)], mitochondrial genomes of mammals from the SibAla region, chloroplast genomes of *Abies sibirica* (NC_035067.1) and *Pinus pumila* (NC_041108.1), genomes of extinct mammals [*Mammuthus primigenius*, TaxID: 37349, (*64*); *Coelodonta antiquitatis*, TaxID: 222863, (*65*)], PhyloNorway (*8*), and viral genomes from IMG/VR v4 (*66*). The reference index was built using Bowtie2 (v2.5.1) with default settings (*61*). Merged reads were aligned against this indexed reference with up to 1000 valid and unique alignments per read (-k 1000). Resulting candidates were then sorted and taxonomically classified using ngsLCA with a minimum identity threshold of 95% (*62*). The full bioinformatic pipeline is available on Zenodo (*20*).

A total of 272,448,770 reads were taxonomically classified. Read counts for each FASTQ file after the major bioinformatic processing steps are summarized in data S2. Five of the 69 controls yielded no classified reads, leaving 64 controls in the taxonomically classified dataset. The ages of the sediment samples ranged from −60 to 58,200 cal yr BP.
